# SARS-CoV-2 expresses a microRNA-like small RNA able to selectively repress host genes

**DOI:** 10.1101/2021.09.08.459464

**Authors:** Paulina Pawlica, Therese A. Yario, Sylvia White, Jianhui Wang, Walter N. Moss, Pei Hui, Joseph M. Vinetz, Joan A. Steitz

## Abstract

Severe acute respiratory syndrome coronavirus 2 (SARS-CoV-2), the causative agent of coronavirus disease (COVID-19), continues to be a pressing health concern. In this study, we investigated the impact of SARS-CoV-2 infection on host microRNA (miRNA) populations in three human lung-derived cell lines, as well as in nasopharyngeal swabs from SARS-CoV-2 infected individuals. We did not detect any major and consistent differences in host miRNA levels after SARS-CoV-2 infection. However, we unexpectedly discovered a viral miRNA-like small RNA, named vmiR-5p (for viral miRNA), derived from the SARS-CoV-2 ORF7a transcript. Its abundance ranges from low to moderate as compared to host miRNAs. vmiR-5p functionally associates with Argonaute proteins — core components of the RNA interference pathway — leading to downregulation of host transcripts. One such host messenger RNA encodes Basic Leucine Zipper ATF-Like Transcription Factor 2 (BATF2), which is linked to interferon signaling. We demonstrate that vmiR-5p production relies on cellular machinery, yet is independent of Drosha protein, and is enhanced by the presence of a strong and evolutionarily conserved hairpin formed within the ORF7a sequence.

**Significance statement:** We discovered that severe acute respiratory syndrome coronavirus 2 (SARS-CoV-2) expresses a small viral non-coding RNA, named vmiR-5p (for viral miRNA), derived from the ORF7a transcript. vmiR-5p associates with the cellular RNA interference machinery to regulate host transcripts likely via target silencing. The production of vmiR-5p relies on cellular machinery and the formation of a strong hairpin within ORF7a sequences. This newly-described vmiR-5p may contribute to SARS-CoV-2 pathogenesis and could become a target for therapeutic intervention.

## Introduction

Coronaviruses are large single-stranded positive-sense RNA viruses that infect various animals; in humans coronaviral infection results in mild to severe respiratory disease. β-coronaviruses, such as severe acute respiratory syndrome coronavirus (SARS-CoV) and Middle East respiratory syndrome coronavirus (MERS-CoV), cause very severe disease in humans, while the recently emerged severe acute respiratory syndrome coronavirus 2 (SARS-CoV-2) results in coronavirus disease (COVID-19) characterized by wide array of symptoms from mild to serious illness. SARS-CoV-2 is the cause of the COVID-19 pandemic due to its high transmissibility and emergence of novel variants (1). It is crucial to understand all aspects of SARS-CoV-2 pathogenesis in order to develop multiple tools to combat the constantly-adapting virus.

microRNAs (miRNAs) are ∼22 nucleotide (nt) small non-coding RNAs (ncRNAs) involved in post-transcriptional regulation of gene expression in animals and plants (2). During canonical miRNA biogenesis, RNA polymerase II transcribes primary miRNAs (pri-miRNAs) characterized by formation of strong hairpins that are recognized and cleaved by the endonuclease Drosha, resulting in ∼70 nt-long precursor miRNAs (pre-miRNAs). Pre-miRNAs are exported to the cytoplasm, where they are subjected to yet another cleavage by Dicer, giving rise to ∼22 nt-long miRNA duplexes, which are then loaded onto Argonaute (Ago) proteins. The passenger strand is removed, and the mature miRNA guides Ago proteins to partially complementary target sequences, usually located in the 3′ untranslated region (UTR) of messenger RNAs (mRNAs). miRNA binding leads to translation inhibition and/or mRNA decay by decapping and deadenylation. In the special case where the miRNA is nearly perfectly complementary to its target mRNA, it directs Ago2 (one of four Ago proteins in humans) to cleave the mRNA target; a function typically associated with small interfering RNAs (siRNAs) (2). These modes of target repression belong to RNA interference (RNAi) pathway, but Ago2-mediated cleavage is more potent in target silencing and requires lower intracellular copy numbers than the canonical miRNA action mode (3). Importantly, miRNA profiles differ between cell types and developmental stages, and a single miRNA can downregulate multiple transcripts, precisely regulating gene expression dependent on the cell’s needs. In humans, most mRNAs are regulated by miRNAs, and aberrant miRNA levels are linked to disease (2).

Viruses often hijack the miRNA pathway; either by depleting host miRNAs or by producing their own miRNAs (reviewed in (4, 5)). Some examples of host miRNA regulation include: selective host miRNA decay mediated by certain herpesviral transcripts in a process called target-directed miRNA degradation (TDMD; reviewed in (6, 7)), and widespread miRNA polyadenylation by poxvirus poly(A) polymerase, which results in miRNA decay (8). Viruses can produce their own miRNAs, often via non-canonical biogenesis pathways; especially cytoplasmic viruses have devised multiple strategies to bypass the requirement for nuclear Drosha (4, 9). It has been proposed that coronaviruses, SARS-CoV and SARS-CoV-2, produce small viral RNAs (svRNAs) involved in viral pathogenesis independently of RNAi pathway (10, 11). One recent study suggested the existence of miRNA-like RNAs produced by SARS-CoV-2, which might contribute to inflammation and type I interferon (IFN) signaling (12).

In this study, by using a small RNA-seq library preparation protocol that selectively enriches for miRNAs, we unexpectedly discovered a miRNA-like viral ncRNA, named vmiR-5p (for viral miRNA), expressed by SARS-CoV-2. We show that it associates with Ago proteins and has the potential to regulate host transcripts likely via target silencing. We demonstrate that vmiR-5p production relies on cellular machinery, but is independent of Drosha proteins and is enhanced by the presence of a strong hairpin formed within ORF7a sequences. This newly-described vmiR-5p may contribute to SARS-CoV-2 pathogenesis and could become a target for therapeutic intervention.

## Results

### SARS-CoV-2 infection has minimal impact on the miRNA population of its host cell

To explore how SARS-CoV-2 affects host miRNA populations, we infected three lung-derived human cell lines either at high multiplicity of infection (MOI of 5) to detect the possible direct impact of the incoming virions on host miRNAs, or at low MOI (0.05) to assess the effect of viral replication on these small RNA species. We used non-small-cell lung cancer cells Calu-3, lung adenocarcinoma cells A549 transduced with the human viral receptor, angiotensin-converting enzyme 2 (ACE2), and lung adenocarcinoma cells PC-9, all of which we found to support SARS-CoV-2 replication (red, yellow and blue dots, respectively, in Fig. 1A). SARS-CoV-2 replicated better in Calu-3 and PC-9 cell lines than in the A549-hACE2 cell line, possibly because the viral receptor is greatly overexpressed in the last cell line (Fig. S1A), which could sequester budding virions at the cell surface.

**Figure 1.**
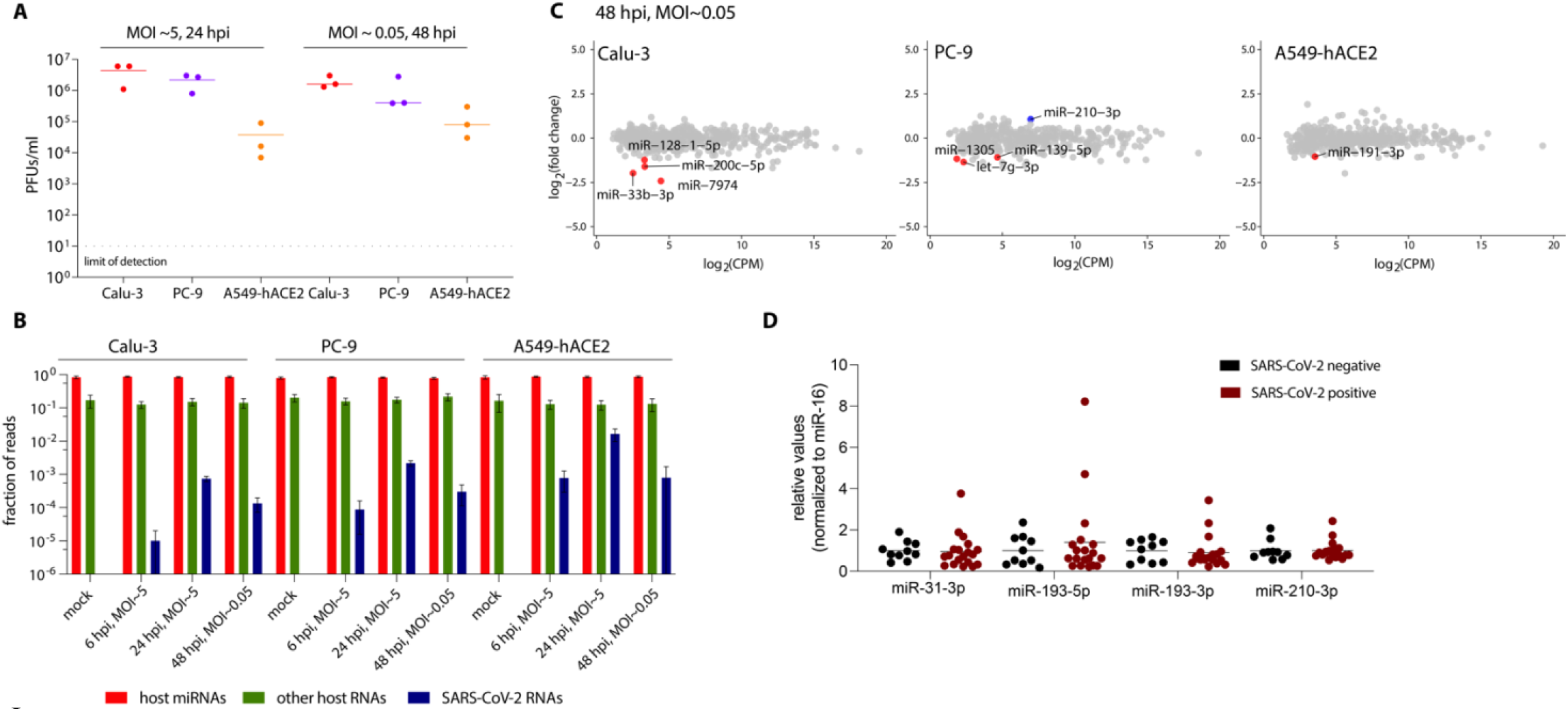
SARS-CoV-2 infection has minimal impact on host miRNA levels. **A)** SARS-CoV-2 replicates in the three human lung cell lines used in this study as demonstrated by plaque assays. **B)** Small RNA library preparation strongly enriches host miRNAs. A bar graph summarizing the origin of obtained reads across the sequenced conditions. Error bars signify SD. **C)** Plots summarizing the fold-change in host miRNA levels 48 hpi with SARS-CoV-2, n = 3. Colored dots denote miRNAs that significantly (p < 0.05) changed (at least two-fold). Significance was calculated with edgeR. **D)** The relative levels of selected miRNAs are unchanged in nasopharyngeal samples from SARS-CoV-2-infected patients versus uninfected controls. PFUs — plaque forming units; hpi — hours post infection; MOI — multiplicity of infection; CPM — counts per million.

Small RNA sequencing was performed at both 6 h and 24 h post infection (hpi) at high MOI and 48 hpi at low MOI, as well as for the uninfected controls (at the 24-h time point). The vast majority (84% ± 5%) of obtained reads mapped to host miRNAs (green bars in Fig. 1B) with an average read length of 22-23 nts (Fig. S1B). Overall, only a small fraction of small RNA reads mapped to the viral genome (0.07±0.01%, 0.21±0.04% and 1.63±0.67% at high MOIs 24 hpi for Calu-3, PC-9 and A549-hACE2, respectively; blue bars in Fig. 1B). This number is in agreement with similar studies of svRNAs in other RNA viruses (10-12).

SARS-CoV-2 infection resulted in minimal alteration of some host miRNA levels, but no miRNA displayed a uniformly significant change across all three cell lines (red and blue dots in Figs. 1C and S1C). To identify any consistent shifts in host miRNA levels across the three cell lines, we generated a heat map of fold-changes for miRNAs that were significantly altered, by at least two-fold, in at least one condition (Fig. S1D). Some of these trends were further examined by Northern blotting of RNA from Calu-3 cells infected with SARS-CoV-2 at low MOI. The most reproducibly changed (upregulated) miRNA, although not reaching statistical significance, was miR-210-3p (Fig. S1E). Interestingly, GO term analyses for validated miR-210-3p targets (13) revealed that this miRNA is involved in regulating responses to cellular stimuli, such as hypoxia (Fig. S1F). In addition, in two datasets obtained from lung biopsies of COVID-19 patients (14, 15) miR-210-3p targets were downregulated to a greater extent than other human mRNAs (Fig. S1G).

Of note, the miRNA changes observed by small RNA-seq were not detected by TaqMan RT-qPCR from nasopharyngeal swabs from SARS-CoV-2-positive individuals (Fig. 1D). Interestingly, analysis of unnormalized Ct values revealed that after SARS-CoV-2 infection the overall miRNA abundance increased, while the levels of ncRNAs from two other small RNA classes (small nuclear RNA U6B and small nucleolar RNA U44) decreased (Fig. S1H), suggesting that miRNAs might escape the viral host shut-off effect (16). This agrees with the reported stabilization of miRNAs in circulating exosomes (17, 18). Thus, it may be possible to devise small RNA-mediated treatments to treat COVID-19. Overall, we did not find any major impact of SARS-CoV-2 infection on host-cell miRNA populations.

### SARS-CoV-2 expresses a miRNA-like small RNA derived from the ORF7a sequence

Of the reads derived from SARS-CoV-2, 5.2% ± 2.8% mapped to a single peak within the ORF7a sequence (Figs. 2A and S2A). ORF7a encodes a SARS-CoV-2 accessory protein, a putative type-I transmembrane protein involved in antagonizing the type I IFN response (reviewed in (19)). The reads mapping to the ORF7a sequence are approximately 20 nt-long (Fig. 2B). Because the small RNA libraries were dominated by host miRNAs and a U (the preferred 5′-terminal nucleotide for Ago loading (20)) is present at the 5′ end, we interrogated whether this RNA could represent a viral miRNA-like ncRNA.

**Figure 2.**
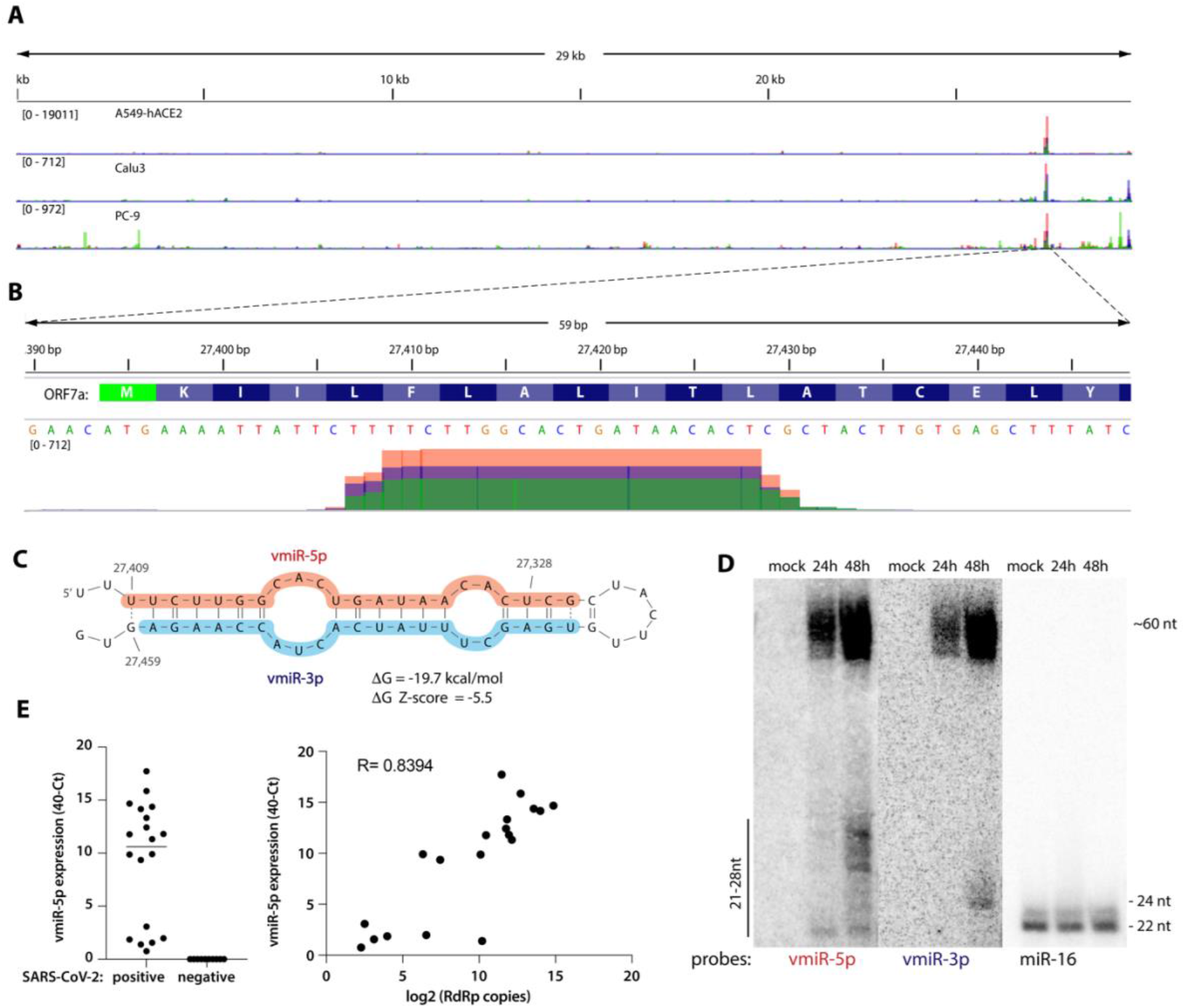
SARS-CoV-2 expresses a small RNA derived from the ORF7a sequence. **A)** Viral small RNA reads obtained from the three cell lines infected with SARS-CoV-2 map to a single distinct peak within the viral genome (data for MOI 5, 24 hpi are shown). The replicates for each cell line were overlayed on a single track (represented by different colors) and are normalized to 10^7^ total reads. MOI — multiplicity of infection. **B)** The reads coming from SARS-CoV-2 (∼20 nt-long) map near the beginning of the ORF7a gene (encoded amino acids are shown above the nucleotide sequence). Data from Calu-3 cells are shown. **C)** vmiR-5p forms a hairpin with the sequence immediately downstream in the viral genome. Shaded nucleotides indicate the sequences detected by Northern blot probes: pink for 5p and blue for 3p. **D)** vmiR-5p can be detected by Northern blotting of extracts from Calu-3 cells infected with SARS-CoV-2 at MOI 0.05. **E)** As measured by custom TaqMan RT-qPCR, vmiR-5p is present in nasopharyngeal samples from SARS-CoV-2-infected individuals (right panel), with its levels correlating with viral load (left panel). RdRp — RNA-dependent RNA polymerase.

Since miRNA biogenesis relies on formation of RNA hairpins, we asked whether such an RNA structure could be found within the viral ORF7a sequence. Indeed, the putative ncRNA together with its downstream region forms a strong hairpin (ΔG = -19.7 kcal/mol; Fig. 2C); this stability value is found to be unusually low given the AU-rich nucleotide composition, suggesting that it arose through an evolutionarily-driven process; indeed, the thermodynamic Z-score was -5.5 (meaning that the WT sequence is more than 5 standard deviations more stable than random). In addition, a comprehensive analysis of RNA structural motifs in SARS-CoV-2 found that this hairpin is one of the few (nine) that show evidence of statistically significant sequence covariation — a feature of evolutionary conservation (21). The formation of the hairpin is also in agreement with previously-reported structure probing reactivity datasets (22-25), where the only reactive nucleotides occur in loop regions. Since the newly-identified ncRNA is derived from this hairpin — which is a hallmark of miRNA biogenesis (26) — we named it viral miRNA-5p (vmiR-5p). There exists sequence and structure similarity of the RNA hairpin between SARS-CoV-2, bat coronavirus RaTG13, and a pangolin coronavirus (Fig. S2B).

vmiR-5p and its pre-miRNA-sized precursor are detectable by Northern blotting of RNA from Calu-3 cells infected with SARS-CoV-2 at low MOI (Fig. 2D). Interestingly, the most prevalent species of vmiR-5p observed by Northern blotting are 25-27 nt-long, longer than those identified by small RNA sequencing. Since the small RNA library was size-selected, this could mean that vmiR-5p is in fact more abundant than when quantified by RNA-seq. However, because miRNAs are usually ∼22 nt-long, vmiR-5p or its precursor could have an additional RNAi-independent function. vmiR-5p was detected in individuals infected with SARS-CoV-2 (Fig. 2E) and its abundance correlated positively with viral load/genomic RNA (as measured by the levels of RNA-dependent RNA polymerase; RdRp).

### vmiR-5p binds Ago to repress human mRNAs

Since the miRNA program is executed by Argonaute proteins, we asked whether vmiR-5p associates with these host proteins. We performed anti-pan-Ago RNA immunoprecipitation (RIP) on extracts from Calu-3 and A549-hACE2 cells infected with SARS-CoV-2 at high MOI, followed by small RNA sequencing. In both cell lines, vmiR-5p associates with Ago proteins, although slightly more weakly than most host-cell miRNAs; its expression levels are comparable to low or moderately expressed host miRNAs (Fig. 3A). To obtain enough material for Northern blotting experiments, we performed an anti-HA RIP from Calu-3 cells transduced with either empty vector or FLAG-HA-tagged human Ago2 (27). Although weak, Ago2 selectively bound vmiR-5p, but not vmiR-3p. In these experiments, we observed slight RNA degradation likely arising during the SARS-CoV-2 inactivation step (30 min incubation at RT). Perhaps this is the reason why vmiR-5p bands in Fig. 3B migrate somewhat faster than those in Fig. 2D.

**Figure 3.**
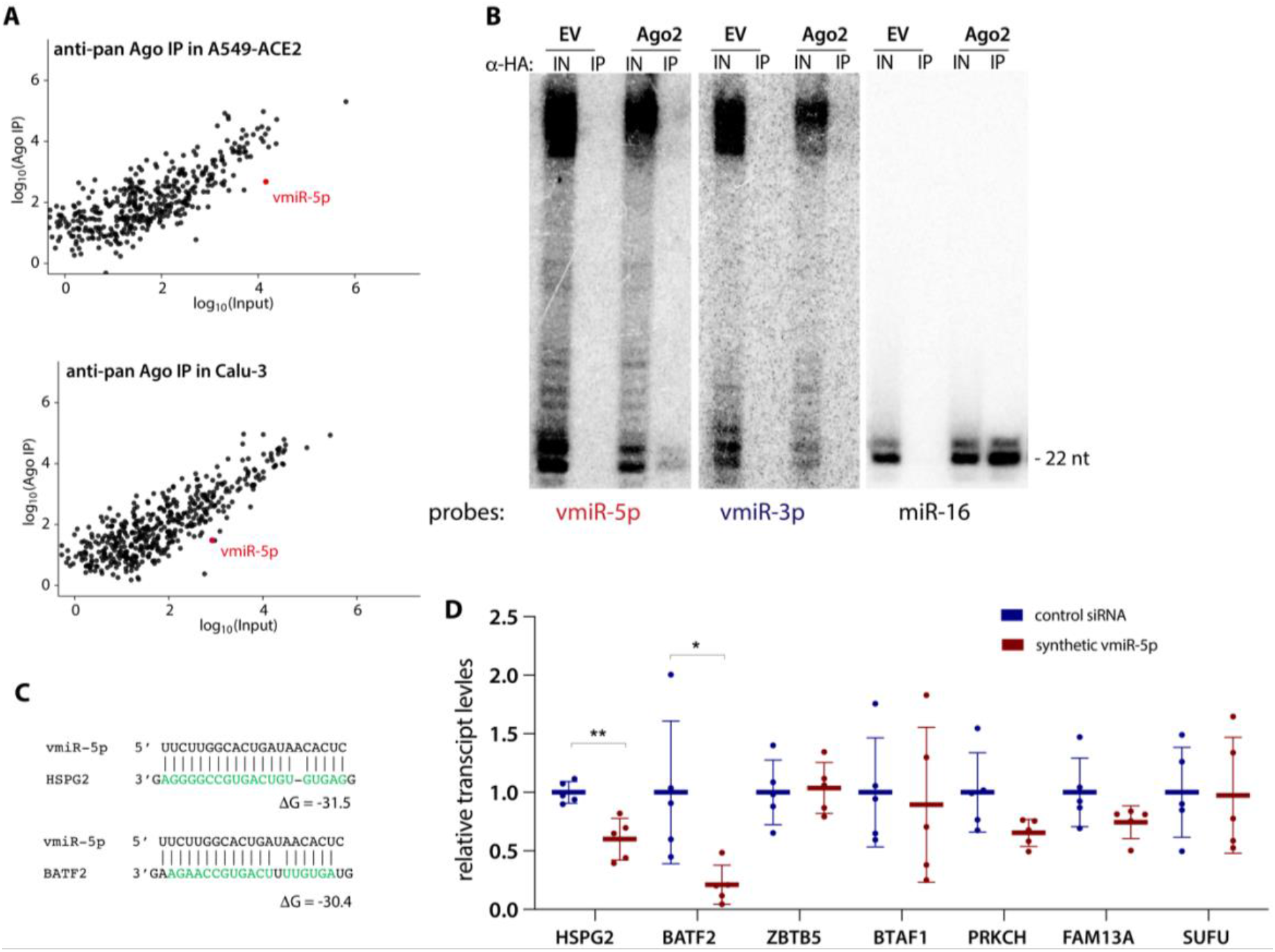
vmiR-5p associates with Argonaute and has the capacity to repress the levels of host transcripts. **A)** vmiR-5p associates with Ago proteins and its levels are comparable to those of moderately expressed host miRNAs. Anti-pan-Ago RNA Immunoprecipitation followed by sequencing was performed on extracts of Calu-3 and A549-hACE2 cells infected with SARS-CoV-2 at MOI 5 for 24h. Each plot shows the average of three independent experiments. **B)** vmiR-5p is selectively loaded on Ago2. Representative Northern blots for viRNA5p and viRNA3p showing anti-HA IP from Calu-3 cells transduced with either empty vector (EV) or with FLAG-HA-tagged Ago2 (Ago2). IN — input, 10%. The miR-16 lanes provide size markers. **C)** Predicted interactions between vmiR-5p and sequences from the CDSs of two targeted host mRNAs. **D)** Synthetic vmiR-5p downregulates host gene expression. HEK293T cells were transfected with synthetic vmiR-5p (annealed to a passenger strand) and levels of seven selected host transcripts shown in Table 1 were measured by RT-qPCR 24 h later. Means with SD are shown. * – p < 0.05 and ** – p < 0.01 as calculated by two-tailed paired t-test.

The role of Ago-associated small RNAs is to repress targeted mRNAs via sequence complementarity, we thus computationally searched the repository of human mRNA transcripts (obtained from GENECODE) for putative vmiR-5p targets. Considering the moderate expression levels of vmiR-5p as compared to those of host miRNAs, we hypothesized that vmiR-5p might rely on Ago2’s ability to cleave its mRNA target. Since Ago2-mediated target cleavage tolerates a limited number of mismatches (28, 29), we searched for mRNAs with high complementarity to vmiR-5p (Table 1; Fig. 3C), and selected some of these for further validation.

**Table 1.**
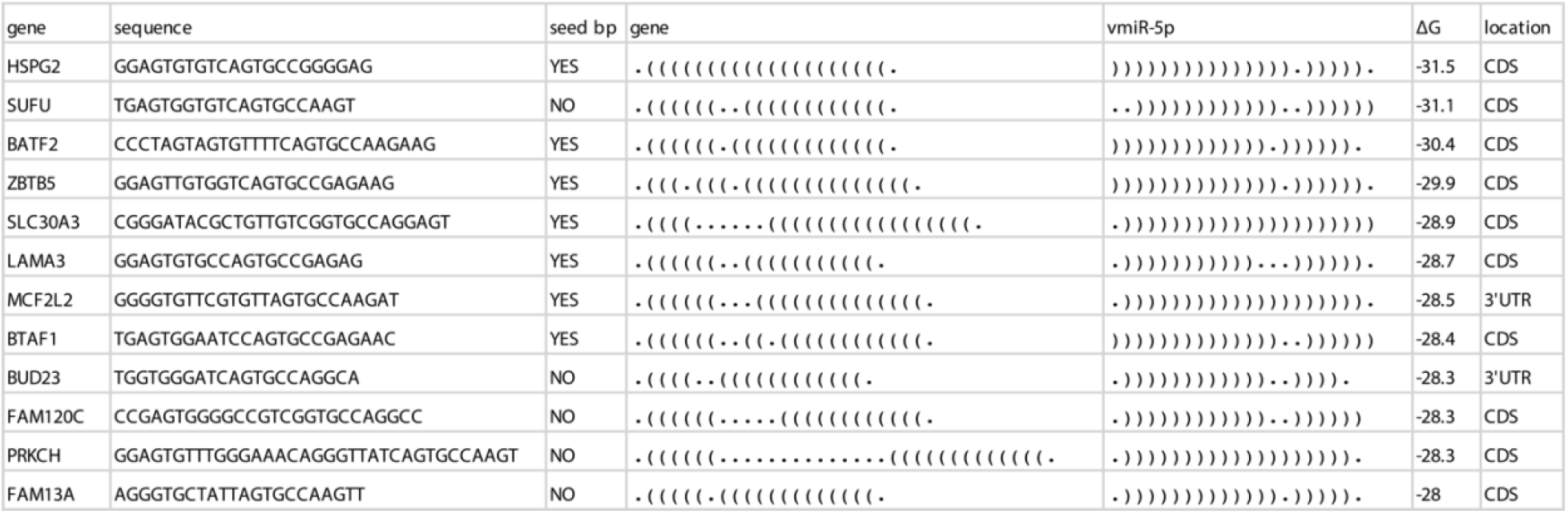
Identification of human transcripts highly complementary to vmiR-5p. bp — base pairing.

Due to the widespread host-shutoff effect after infection with SARS-CoV-2 (16), we were unable to perform meaningful quantification of mRNAs from infected cells. We thus transfected synthetic vmiR-5p (annealed to a passenger strand) into HEK293T cells and assayed transcript levels of predicted target mRNAs (Fig. 3D). Two mRNAs, Basic Leucine Zipper ATF-Like Transcription Factor 2 (BATF2) and (Heparan Sulfate Proteoglycan 2) HSPG2, were significantly downregulated (Fig. 3D). BATF2 has been linked to INF-gamma signaling via association with Irf1 (30). HSPG2 is the core protein of a large multidomain proteoglycan that binds and cross-links many extracellular matrix components; it interestingly participates in the attachment of some viruses, including the coronavirus NL63 (31). We hypothesize that vmiR-5p has the potential to repress human mRNAs that support the host IFN response, and perhaps to inhibit viral superinfection.

### Production of functional vmiR-5p depends on cellular machinery

To assess whether vmiR-5p biogenesis can occur in a virus-free system, we transfected *in vitro* transcribed RNA into HEK293T cells. A 100 nt-long viral RNA sequence (WT), alongside counterparts containing mutations either in the 3′ portion (m3p) or in the 5′ portion (m5p) of the hairpin (Fig. 4A) were transfected into cells and RNA samples collected after 6 h were analyzed by Northern blotting (Fig. 4B). The WT sequence was processed inside cells to produce a 21 nt-long band of vmiR-5p, while the m3p did not yield efficient production of an RNA of such size.

**Figure 4.**
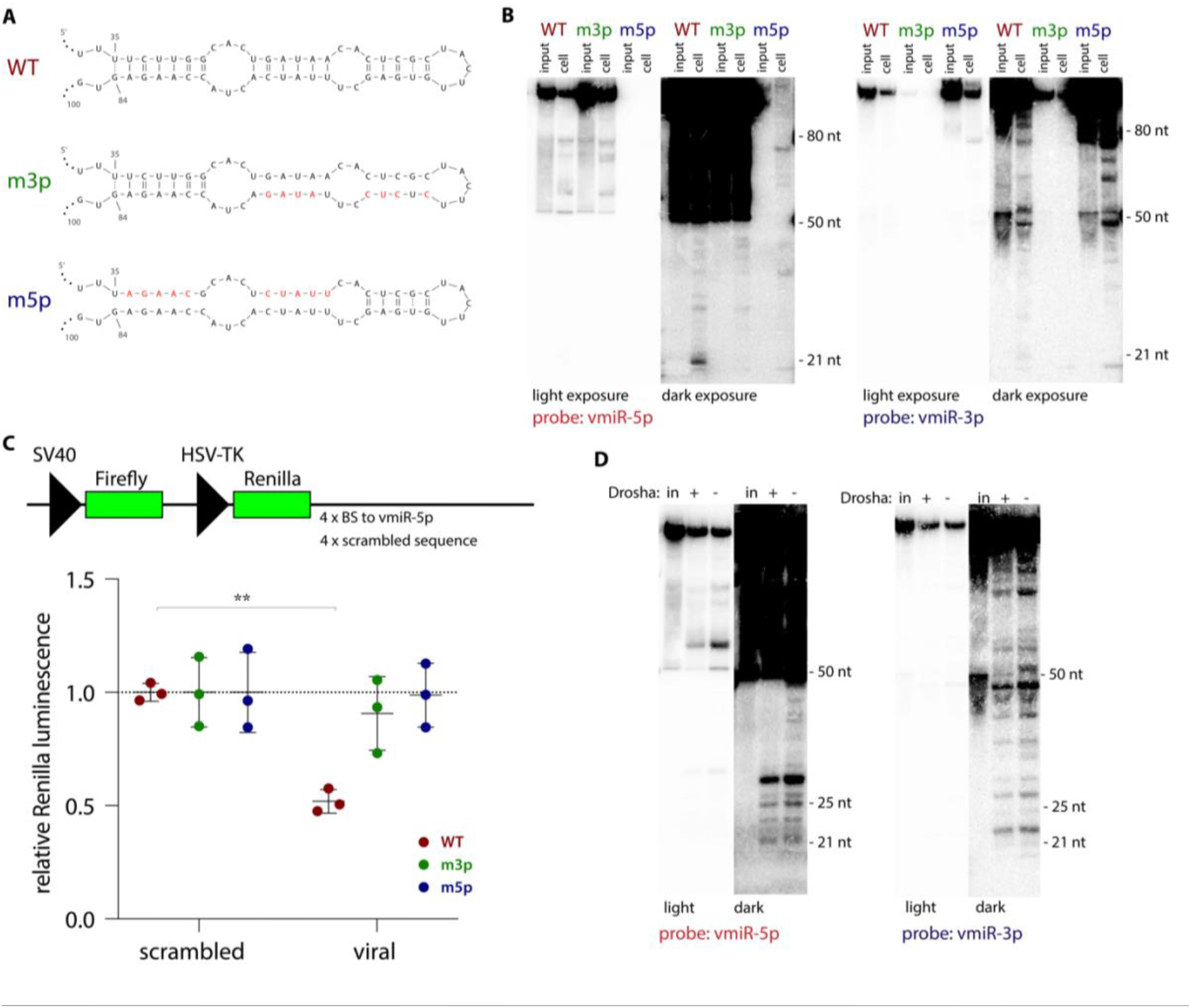
Production of functional vmiR-5p is Drosha-independent. **A)** *In vitro* transcripts used in this study. Red nts represent introduced mutations. **B)** vmiR-5p processing inside cells is enhanced by the formation of a hairpin involving the downstream sequence. A representative Northern blot of RNAs isolated from HEK293T cell 6 h post transfection with each of the *in vitro* transcribed 100 nt-long RNA sequences shown in panel A. Input, 10%. **C)** vmiR-5p processed inside HEK293 cells is capable of repressing a reporter. Top panel: schematic of the psiCHECK-based luciferase reporters used. Each of the four binding sites (BS) was 21 nt-long, separated by unique 20 nt-long spacers. Bottom panel: vmiR-5p generated from an *in vitro*-transcribed hairpin can regulate a luciferase reporter gene containing downstream complementary sites in HEK293T cells. Means with SD are shown. ** – p < 0.01 as calculated by two-tailed paired t-test. **D)** vmiR-5p processing is Drosha independent. A representative Northern blot of RNAs isolated from either Huh7.5 or Huh7.5 Drosha knockout cells (32) 6 h post transfection with *in vitro*-transcribed 100 nt-long viral RNA sequence. in — input, 10%.

To investigate whether the produced RNA functionally associates with Ago proteins, we transfected *in vitro* transcribed RNA together with luciferase reporters containing 3′ UTR sequences complementary to either vmiR-5p or scrambled controls. The luciferase activity of the reporter containing viral sequences was significantly decreased in the presence of the WT ORF7a RNA (Fig. 4C), but not by the other constructs. Finally, we used Drosha knockout cell lines (32) to show that the production of vmiR-5p occurs independently of this enzyme’s activity (Fig. 4D). These results clearly demonstrate that vmiR-5p is processed from a hairpin independently of viral proteins and of cellular Drosha, and that it can be functionally loaded on Ago.

## Discussion

Here, we began by investigating the impact of SARS-CoV-2 infection on host miRNA populations using three human lung-derived cell lines (at various MOIs) and found no consistent changes detected in all three cell lines (Figs. 1C and S1C). We also assessed the levels of selected miRNAs in patient samples and again did not detect major differences after SARS-CoV-2 infection (Fig. 1D). Other groups have also ventured into examining miRNA profiles during SARS-CoV-2 infection, but there is little overlap between these prior studies (33-35) and our results. Differences between the outcomes likely stem from the use of different methods of RNA-seq library preparation (36, 37). Our libraries were prepared with the NEXTflex v2 kit, which has been shown to be highly selective for miRNAs (36), and contained ∼84% host miRNAs (Fig. 1B), while libraries that rely on template switching used by others (33) yield only ∼17% miRNAs (37). The presence of additional reads in the latter libraries could result in nonspecific alignment to miRNA loci. For example, miR-155-3p, which is believed to represent a host miRNA passenger stand, was reported to increase upon SARS-CoV and SARS-CoV-2 infection (33), but we were unable to detect this miRNA either by sequencing or by Northern blotting (data not shown). Similarly, miR-4485 was described to be upregulated in another study (34), but it did not pass the abundance threshold (> 1 CPM) in our small RNA-seq analysis. Also, other studies assessing the impact of various coronaviruses on host miRNA levels do not support the notion that coronaviruses have major impact on host miRNA populations (38-41). We cannot rigorously exclude the possibility that some host miRNAs levels are altered during SARS-CoV-2 infection, but if so, these changes are likely minor.

Interestingly, in samples from individuals infected with SARS-CoV-2 all assessed host miRNAs seemed to be stabilized, while — similar to what is known about host mRNAs in infected cells (16) — U6B snRNA and U44 snoRNA levels decreased (Fig. S1H). These results suggest that miRNAs might be resistant to the viral host-shutoff effect. Indeed, miRNAs can be very stable because of their association with Ago proteins, especially when secreted in exosomes (2, 17, 18). It is tempting to speculate that, because of their high stability, small RNAs could be successfully used as therapeutic agents against SARS-CoV-2. Regardless, we caution against examining miRNA profiles using RT-qPCR and normalizing to ncRNAs from different classes, such as snRNAs and snoRNAs.

In this study, we discovered that SARS-CoV-2 expresses an miRNA-like small RNA, which we call vmiR-5p (Fig. 2). There are multiple examples of svRNAs expressed by RNA viruses, such as influenza (42), enterovirus 71 (43), Hepatitis C (44), hepatitis A (45), Polio (44), Dengue (44, 46), Vesicular Stomatitis (44), West Nile (44, 47), coronaviruses (10-12) and retroviruses (48, 49). It has been suggested that some svRNAs function through the host-cell RNAi pathway (12, 44, 45), but most proposed roles are independent of this machinery (10, 11, 42, 43, 50). As compared to other studies of SARS-CoV-2, we did not detect svRNAs derived from the N gene (10, 12), which suggests that the functions of those is not related to RNAi pathway. Interestingly, vmiR-5p was also detected by Meng and colleagues, but was not selected for further validation (12).

vmiR-5p is derived from the beginning of the coding sequence of the ORF7a transcript (Fig. 2B), which is the most abundant subgenomic viral RNA detected during SARS-CoV-2 infection (33, 51). The sequence of vmiR-5p is preserved in all SARS-CoV-2 isolates and variants of concern, and is related to those of two other coronaviruses (Fig. S2). Interestingly, ORF7a deletions, which lead to decreased innate immune suppression, occur frequently in SARS-CoV-2, but affect almost exclusively the C-terminus of the protein, while the N-terminus — where the vmiR-5p sequence is located — is preserved (52). These data and the observed conservation of the hairpin containing vmiR-5p (21) support the notion that this sequence is functional.

The abundance of vmiR-5p ranges from low to moderate as compared to host miRNAs (Fig. 3A), similar to other svRNAs (10, 45, 53), and is consistent with the idea that its functionality relies on high complementarity to target mRNA, likely inducing mRNA cleavage. Indeed, we have shown that vmiR-5p associates with human Ago proteins and that it can repress human targets (Fig. 3). Using bioinformatics, we identified two host transcripts that are significantly downregulated in the presence of a synthetic vmiR-5p, BATF2 and HSPG2 (Fig. 3 and Table 1). In addition to these, there are likely more targets that may be identified by cross-linking ligation and sequencing of hybrid (CLASH; (54)) methodology. To address the functionality of vmiR-5p in the context of viral infection, we attempted to use luciferase reporters, but due to the host-shutoff effect there was almost no luciferase signal in infected cells. We avoided using antisense oligonucleotides to block vmiR-5p because they would undoubtedly also suppress production of the ORF7a protein and inhibit viral replication. Upcoming studies will focus on the construction of mutant viruses in which the ORF7a coding sequence and hairpin formation are preserved.

Another intriguing possibility is that SARS-CoV-2 uses vmiR-5p to regulate the level of its own negative-sense subgenomic RNAs, or of antigenomic RNAs (55). It is also plausible that the viral hairpin, in addition to being the source of a viral miRNA, could have an additional function independent of the RNAi pathway. RNA viruses have been shown to express regulatory svRNAs. For example, svRNA1 from enterovirus-71 binds to a viral internal ribosome entry site (IRES) to regulate translation of viral proteins (43). Another example is svRNAs from influenza that associate with viral RdRp, possibly enabling the switch from transcription to replication (50).

Finally, we addressed vmiR-5p biogenesis and were able to demonstrate that its processing occurs via cellular machinery and relies on the formation of an RNA hairpin, but is independent of Drosha protein (Fig. 4). Yet, it is possible that viral genes enhance this processing pathway. Many viruses bypass Drosha to produce their pre-miRNAs, e.g. by utilizing RNase Z (56, 57), Integrator complex (58) or a viral protein (such as HIV-1 TAT) (59). In some instances, viral miRNAs biogenesis even relies on Ago2 cleavage (4). We speculate that Dicer, which is responsible for production of the majority of host and viral miRNAs (2, 45, 59, 60), is involved in vmiR-5p biogenesis. Future efforts will aim to uncover which enzymes are responsible for vmiR-5p production.

In summary, viruses develop multiple ways to suppress host gene expression, and multiple overlapping mechanisms often evolve. SARS-CoV-2 regulates host mRNA expression on many levels: stability (16), export (16, 60), translation (61-63) and splicing (61). It is not surprising that yet another strategy, which utilizes the RNAi pathway, to selectively target host — and perhaps viral — transcripts has evolved.

## Acknowledgements

We thank the members of the Steitz lab for advice and support. We are grateful to Dr. Craig Wilen for the help in setting up BSL3 work. We thank Dr. Charles Rice for Hu7.5 cells and Huh7.5 Drosha knockout cells (32). We thank Dr. Benjamin Israelow for a plasmid expressing hACE2. We acknowledge the hard work of Yale’s Environmental Health and Safety program, in particular Benjamin Fontes. This work was funded by the NIH supplement 3P01CA016038-45S1 (to J.A.S.), by NIH grant R01GM133810 grant (to W.N.M)., and by the Yale School of Medicine Department of Internal Medicine, the Prostate Cancer Foundation, the Harrington Discovery Institute, and the COVID-19 Early Treatment Fund (to J.M.V.). P.P. was supported by NIH fellowship K99/R00 (K99GM129412/ R00GM129412). J.A.S. is an investigator of the Howard Hughes Medical Institute.

## Author Contributions

P.P. and J.A.S. designed experiments and interpreted the results. P.P. performed experiments and analyzed data. T.A.Y. provided technical assistance. S.W., J.W., P.H., J.M.V. provided RNA from nasopharyngeal swabs, W.N.M. contributed to computational analyses, P.P. and J.A.S. wrote the manuscript.

## Materials and Methods

### Cells

Calu-3 (ATCC) cells were cultured in Eagle’s minimal essential medium (EMEM, ATCC) with 10% fetal bovine serum (FBS) and Penicillin/Streptomycin (Pen/Strep; GIBCO). PC-9 cells (kind gift from Dr. Craig Wilen) were cultured in RPMI medium (GIBCO) with 10% FBS and Pen/Strep. A549 (ATCC) cells were transduced as described previously (27) with hACE2 plasmid and cultured in F-12 medium (GIBCO) with 10% FBS, Pen/Strep and 1 μg/ml of puromycin (GIBCO). Vero-E6 (ATCC), Huh7.5 and Huh7.5 Drosha knockout (kind gift from Dr. Charles Rice; (32)) cells were cultured in DMEM with 10% FBS, and Pen/Strep. Calu-3 cells were transduced either with FLAG-HA-Ago2 (27) or pLVX (for empty vector control) and cultured in the presence of 1 μg/ml of puromycin.

### Viral stocks

To generate SARS-CoV-2 stocks, Vero-E6 were inoculated with P2 of SARS-CoV-2 isolate USA-WA1/2020 (kind gift from Dr. Craig Wilen) at 0.01 multiplicity of infection (MOI) for 1 h, after which the inoculum was replaced with DMEM with 5% FBS. After 3 days the supernatant was harvested and clarified by centrifugation, concentrated on Ultra-15 Centrifugal Filters (Amicon), aliquoted and stored at −80°C. Virus titers were determined by plaque assay using Vero-E6 cells. All infectious growth was performed in a Biosafety Level 3 laboratory and was approved by the Yale University Biosafety Committee.

### Plaque assays

Vero-E6 cells were seeded at 4×10^5^ cells/well in 12-well plates. The next day, medium was removed, and cells were incubated for 1 h at 37°C with 100 μl serially diluted sample; plates were rocked every 15 min. Next, 1 ml of the overlay media (DMEM, 2% FBS, 0.6% Avicel) was added to each well. 3 days later, the plates were fixed with 10% formaldehyde for 30 min, stained with crystal violet solution (0.5% crystal violet in 20% methanol) for 30 min, and then rinsed with water to visualize plaques.

### Samples for small RNA sequencing

For small RNA sequencing, 5×10^5^ of Calu-3, 5×10^5^ of PC-9 or 1×10^5^ of A549-hACE2 cells were plated per well in 6-well plates. Cells were infected with SARS-CoV-2 at MOI either of 5 or 0.05 for 1 h in 200 µl of inoculum and incubated for 1 h at 37°C. After that, the cells were rinsed with PBS to remove the unbound virus and fresh were media added. Cells were incubated for either 6 h and 24 h (MOI 5), or 48 h (MOI 0.05). In addition, mock-treated controls were collected after 24h. Cells were harvested by removing the media and adding TRIzol at each time point; supernatants were kept to assess viral titer by plague assay. RNA isolation and library preparation was performed under BSL2+ confinement; RNA concentration was measured by Qubit Fluorimeter (ThermoFisher) inside a biosafety cabinet. 1 μg of total RNA was used for library preparation using NEXTflex v2 kit (PerkinElmer) according to the manufacturer’s instructions. Libraries were amplified for 16 cycles and the cDNA libraries were sequenced on the HiSeq 2500 Illumina platform.

### Small RNA data processing

The reads were trimmed of adaptors using Cutadapt (64) with the following settings -u 4 -O 7 -a N{4}TGGAATTCTCGGGTGCCAAGG -q 10 -m 18 -M. The reads were mapped with bowtie2 (65) (--very-sensitive-local) to an index containing human and SARS-CoV-2 genomes. miRNAs were counted by using featureCounts (66) and annotations obtained from miRbase (67). For Ago IP counts, vmiR-5p was added to the annotation file and treated as a host miRNA. Differential expression was determined using edgeR (68). Track visualization was performed using an IGV browser (69) of generated with BEDtools (70) bed files.

### Patient samples and quantitative RT-PCR

The human study reported here from which nasopharyngeal swab samples were obtained was approved by the Yale Human Research Protection Program, (Protocol 2000027971).

The nasopharyngeal swab samples were extracted using MagMAX mirVana Total RNA Isolation Kit (Catalog # A27828), with the script A27828_FLEX_Biofluids for miRNA extraction from biofluid samples on the KingFisher FLEX-96 Magnetic Particle Processor. Briefly, each 100 μl specimen of nasopharyngeal swab in transportation medium was processed and the miRNA-enriched RNA was collected in 50 μl of elution buffer. miRNA RT-qPCR was performed using TaqMan MicroRNA Assay (Applied Biosystems); for miR-16, miR-210-3p, miR-31-3p, miR-193-5p, miR-193-3p, U6B and U44 pre-designed assays were ordered (Cat # 4427975) and for vmiR-5p, a custom assay to detect the sequence UUCUUGGCACUGAUAACAC was ordered (Cat # 4398987). RT-qPCR was performed according to the manufacturer’s guidelines; briefly, cDNA was prepared separately for each miRNA using the TaqMan MicroRNA Reverse Transcription Kit (Applied Biosystems) and qPCR was performed using the TaqMan Fast Advanced Master Mix (Applied Biosystems). For detection of viral RdRp, cDNA was prepared using random primers and SuperScript III (Invitrogen) and qPCR was performed according to the protocol (71) using oligonucleotides and a probe ordered from IDT (sequences are given in Table 2). All work was done in BSL2+ conditions.

**Table 2.**
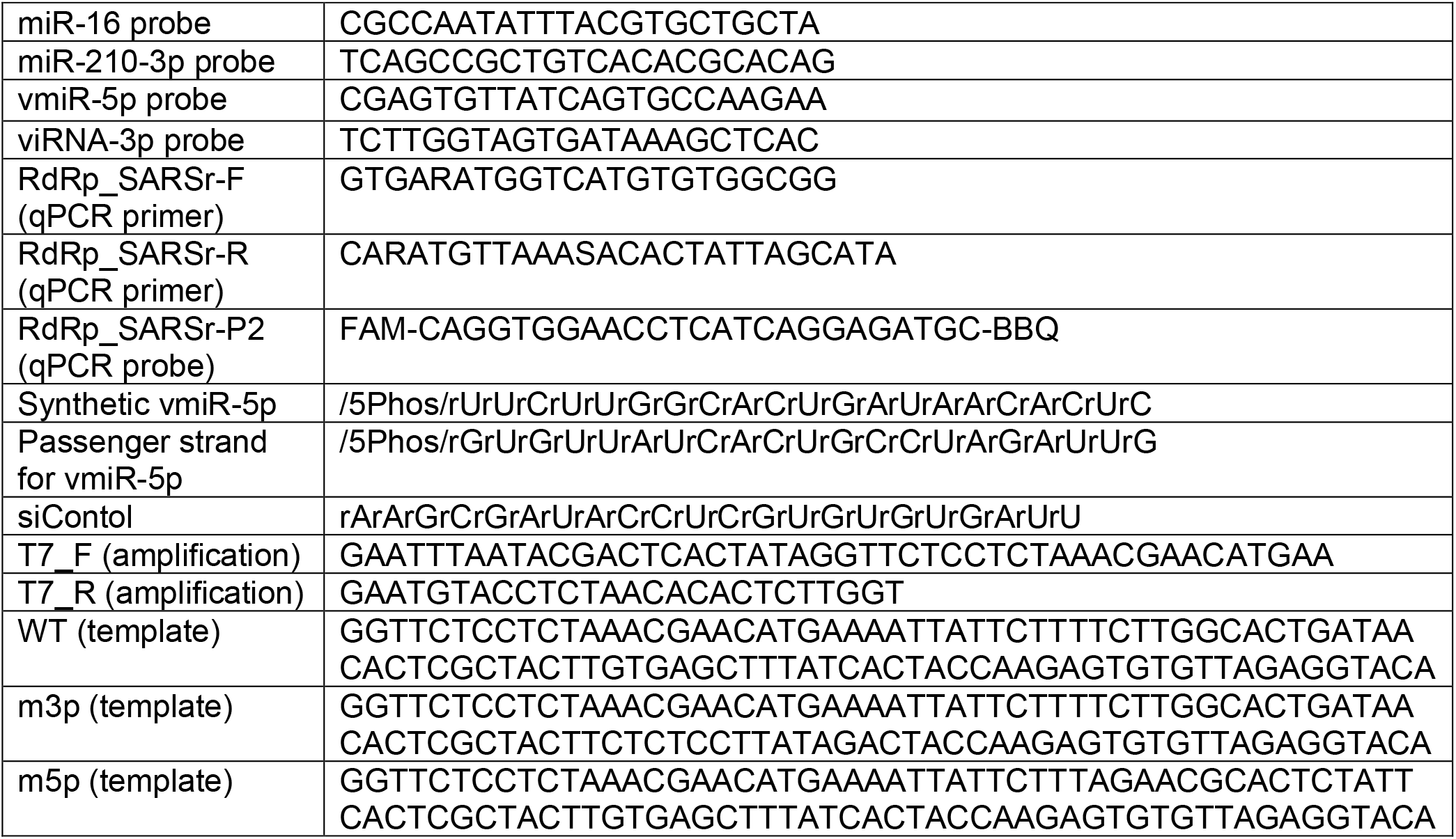
Oligonucleotides used in the study

### Northern blot analysis

5×10^6^ of Calu-3 cells were infected with SARS-CoV-2 at MOI 0.05 in T75 flasks for 1h, after which the inoculum was replaced by fresh media, and cells were incubated for either 24 h or 48 h (mock-treated samples were collected after 48h). Samples were inactivated for 30 min in 5 ml of TRIzol and RNA was isolated in BSL2+ conditions. 10 -15 µg of total RNA was separated by 15% urea-PAGE, electrotransferred to Hybond-NX membrane (Amersham) and crosslinked with 1-ethyl-3-(3-dimethylaminopropyl) carbodiimide (EDC) (72). miRNAs were detected using ^32^P 5’-radiolabeled DNA probes. Densitometry was performed by using Quantity One.

### miR-210-3p target and re-analysis of public data

Counts from RNA-seq experiments of lung biopsies were obtained from (14, 15). The data were divided into miR-210-3p targets (data from miRTarBase (73); excluding “weak” targets) and other transcripts. The data were plotted as cumulative distribution functions and compared to each other using Wilcoxon test.

### Immunoprecipitation

For anti-pan Ago IP, 5×10^5^ of Calu-3 cells or 1×10^5^ of A549-hACE2 cells were infected with SARS-CoV-2 at MOI 5 for 24h. Cells were detached with trypsin-EDTA, pelleted and resuspended in 500 µl of NET-2 buffer (50 mM Tris [pH 7.5], 150 mM NaCl, and 0.05% NP-40). After 30 min of cell lysis inside the BSC, the nuclei were pelleted and supernatants were transferred to 2.0 ml screw-top tubes with O-rings containing magnetic beads (SureBeads Protein G Magnetic Beads, NEB) coupled to antibodies (anti-pan Ago antibody, clone 2A8; Sigma Millipore). 10% of supernatants were kept for input (in TRIzol). The tubes were sealed, decontaminated and placed inside 15 ml screw-top falcon tubes. The 15 ml falcon tubes were decontaminated and transferred to the fridge containing the rotator where the immunoprecipitation took place. After 6h, the beads were washed inside BSC 7 times with NET-2 by gentle pipetting and with using a magnetic stand. Finally, the magnetic beads were resuspended in 1 ml of TRIzol. Described procedures were performed in a Biosafety Level 3 laboratory and were approved by the Yale University Biosafety Committee. RNA isolation and library preparation (as described above) was performed under BSL2+ confinement.

For anti-HA IP, 5×10^6^ of Calu-3 cells, transduced with either EV or Ago2, were infected with SARS-CoV-2 at MOI 0.1 for 48h. Cells were detached with trypsin-EDTA, pelleted and resuspended in 2 ml of Pierce IP Lysis Buffer (Thermo Fisher). Cells were incubated at room temperature for 30 min (approved by the Yale University Biosafety Committee SARS-CoV-2 inactivation method). After this time, the samples were moved to the BSL2+ laboratory, sonicated using a Diagenode Bioruptor Pico sonication device, and IP was performed using Anti-HA Magnetic Beads (Pierce). Beads were resuspended in TRIzol, RNA was extracted and analyzed by Northern blot as described above.

### Western blot analysis

IPs were done as described above (excluding SARS-CoV-2 infection), supernatants were mixed with 4X SDS-PAGE loading buffer. Typically, 25 µl (corresponding to ∼40 µg total protein) were separated on a 10% SDS-PAGE gel, and electrotransferred to a PVDF membrane (BioRad). After blocking with 5s% milk in 1× TBST (20 mM Tris [pH 7.5], 150 mM NaCl, 0.1% Tween 20), the membrane was probed with the appropriate antibodies and detected with Western Lightning Plus-ECL (PerkinElmer) using a Gbox (SYNGENE). Primary antibodies used were anti-FLAG M2 (Sigma Millipore), anti-Ago2 (MA5-23515, Invitrogen) and anti-GAPDH (Cell Signaling).

### Target predictions

Custom Perl scripts that use the RNAduplex algorithm — part of the Vienna RNA Package (53) — were used to hybridize the vmiR-5p sequence to mRNA transcripts obtained from GENCODE (v38) (74) fragmented into 50-nt windows with 5-bp steps. Fragments with the lowest hybridization energy (ΔG) were chosen for further analysis.

### Synthetic RNA transfections and qPCR

Synthetic RNAs (see Table 2 for sequences) were annealed by heating equimolar concentrations for 1 min at 90°C in siRNA buffer (Horizon, 60 mM KCl, 6 mM HEPES-pH 7.5, 0.2 mM MgCl2) and then incubating for 1 h at 37°C. 5×10^5^ HEK293T cells were transfected with 30 μM of either vmiR-5p or control siRNA, by using Lipofectamine RNAiMAX Transfection Reagent (Invitrogen). Cells were collected 48 h later, RNA extractions were performed using TRIzol, samples were treated with RQ1 DNase (Promega). cDNA was made using SuperScript III and random primers (Invitrogen), and qPCR was performed using FastStart Essential DNA Green Master (Roche). Primer sequences are given in Table 2. Results were analyzed using the comparative Ct method (75).

### *In vitro* RNA transcription and processing

The sequences were PCR amplified from templates containing desired mutations and by using flanking primers; the forward primer contained the T7 promoter (see Table 2 for sequences). The product was purified and used for *in vitro* transcription at a final concentration 25 ng/µl. The reaction was carried out for 4 h at 37°C and contained 400 mM HEPES pH 7.5, 120 mM MgCl2, 200 mM DTT, 10 mM Spermidine, 4 mM of each rNTPs, 20 U RNase Inhibitor (Roche), and 5 U of laboratory-made T7 RNA polymerase. The product was purified by 8 M urea 6% polyacrylamide gel electrophoresis (PAGE), extracted in G-50 buffer (20 mM Tris pH 7.5, 0.3 M NaOAc, 2 mM EDTA, 0.1% SDS), and phenol extracted. To ensure hairpin formation, RNAs were annealed by heating in siRNA buffer for 1 min at 90°C and then incubating for 1 h at 37°C.

5×10^5^ cells (HEK293Ts, Huh7.5 or Huh7.5 KO) were transfected with 30 μM of *in vitro* transcribed RNAs, by using Lipofectamine RNAiMAX Transfection Reagent (Invitrogen). Cells were collected 6 h later, RNA extractions were performed using TRIzol and samples were processed for Northern blotting as described above.

### Luciferase reporter assays

4 sites complementary to vmiR-5p (or scrambled sequence as control) separated by 20 nt were PCR amplified using four primers overlapping at the unique spacer sequences (listed in Table 2). The PCR products were cloned downstream of *Renilla* luciferase of psiCHECK(TM)-2 vector (Promega) using XhoI and NotI sites.

5×10^5^ cells (HEK293Ts, Huh7.5 or Huh7.5 KO) were transfected with 30 μM of *in vitro* transcribed RNAs, by using Lipofectamine RNAiMAX Transfection Reagent (Invitrogen). Cells were collected 6 h later, RNA extractions were performed using TRIzol and samples were processed for Northern blotting as described above. 2 4h later, 10 ng psiCHECK reporters and 2 μg pBlueScript II (Stratagene) were transfected using TransIT-293 Transfection Reagent (Mirus). After additional 24h, Firefly and *Renilla* luciferase activities were measured by the Dual-Luciferase Reporter Assay System (Promega) on a GloMax-Multi+ Microplate Multimode Reader (Promega) according to the manufacturer’s instructions.

## Supplementary figures

**Figure S1.**
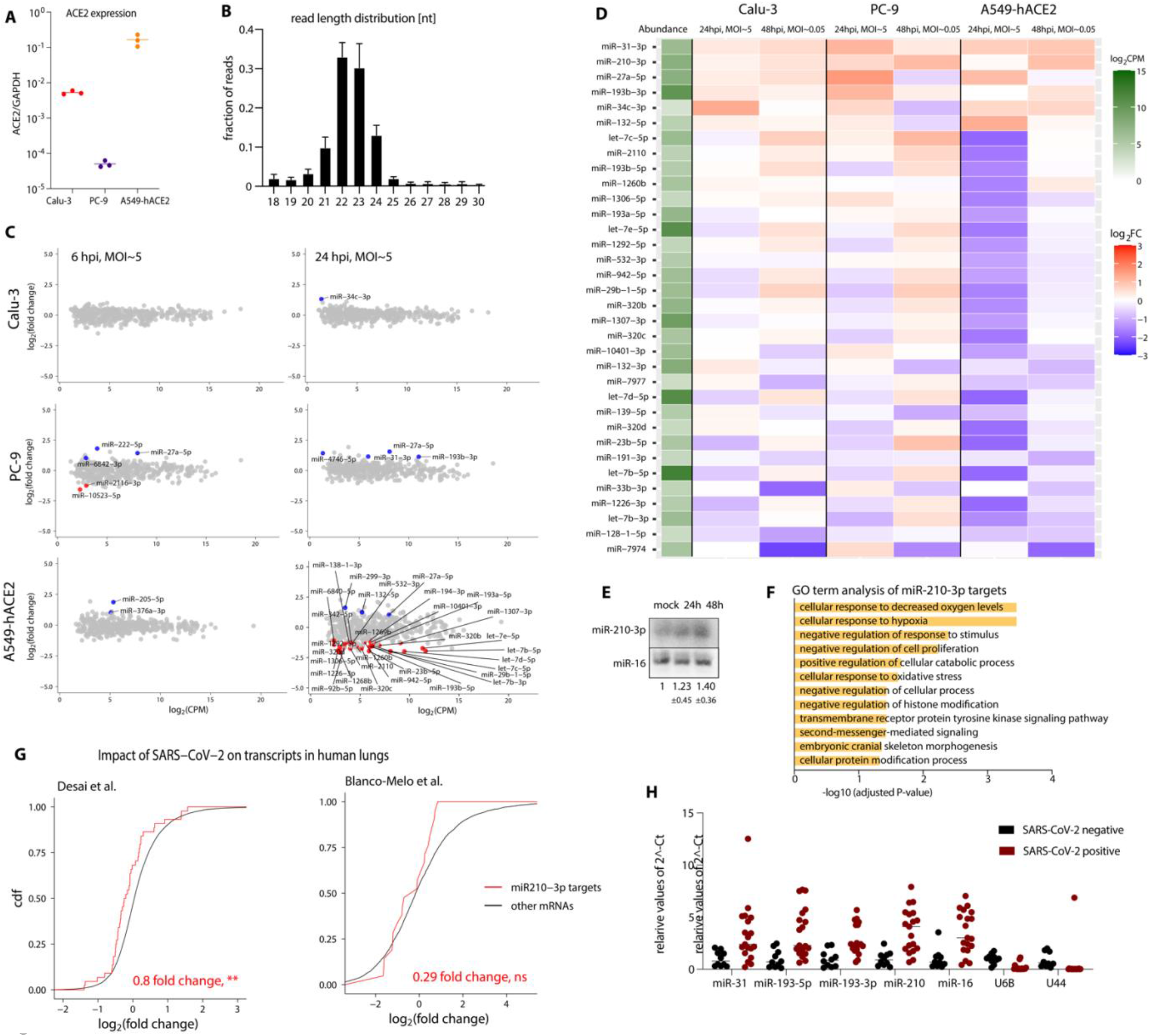
SARS-CoV-2 infection has minimal impact on host miRNA levels. **A)** Relative transcript levels of hACE2 in the three lung cell lines used in the study. **B)** A bar graph summarizing the sizes of obtained reads from all conditions sequenced. Error bars signify SD. **C)** Plots summarizing the fold-change in miRNA levels upon SARS-CoV-2 infection of various cell lines, n = 3. Colored dots denote miRNAs that significantly (p < 0.05) changed (at least two-fold). Significance was calculated using edgeR. **D)** A heat-map showing fold changes (FC) in miRNA levels after SARS-CoV-2 infection. The data are shown only for miRNAs that significantly changed (at least 2-fold) in any of the conditions. **E)** Northern blot showing that miR-210-3p slightly increases after infection with SARS-CoV-2 infection of Calu-3 cells at MOI 0.05, n = 5. **F)** GO term analyses for miR-210-3p targets suggest that this miRNA regulates responses to various stimuli. ** - p = 0.004, as calculated by Wilcoxon test. **G)** miR-210-3p targets (13) exhibit stronger downregulation than other host mRNAs after SARS-CoV-2 infection in human lungs (data replotted from (14, 15)). **H)** Host miRNAs may escape host shutoff induced by SARS-CoV-2 infection. hpi — hours post infection; MOI — multiplicity of infection; CPM — counts per million; cdf — cumulative distribution function.

**Figure S2.**
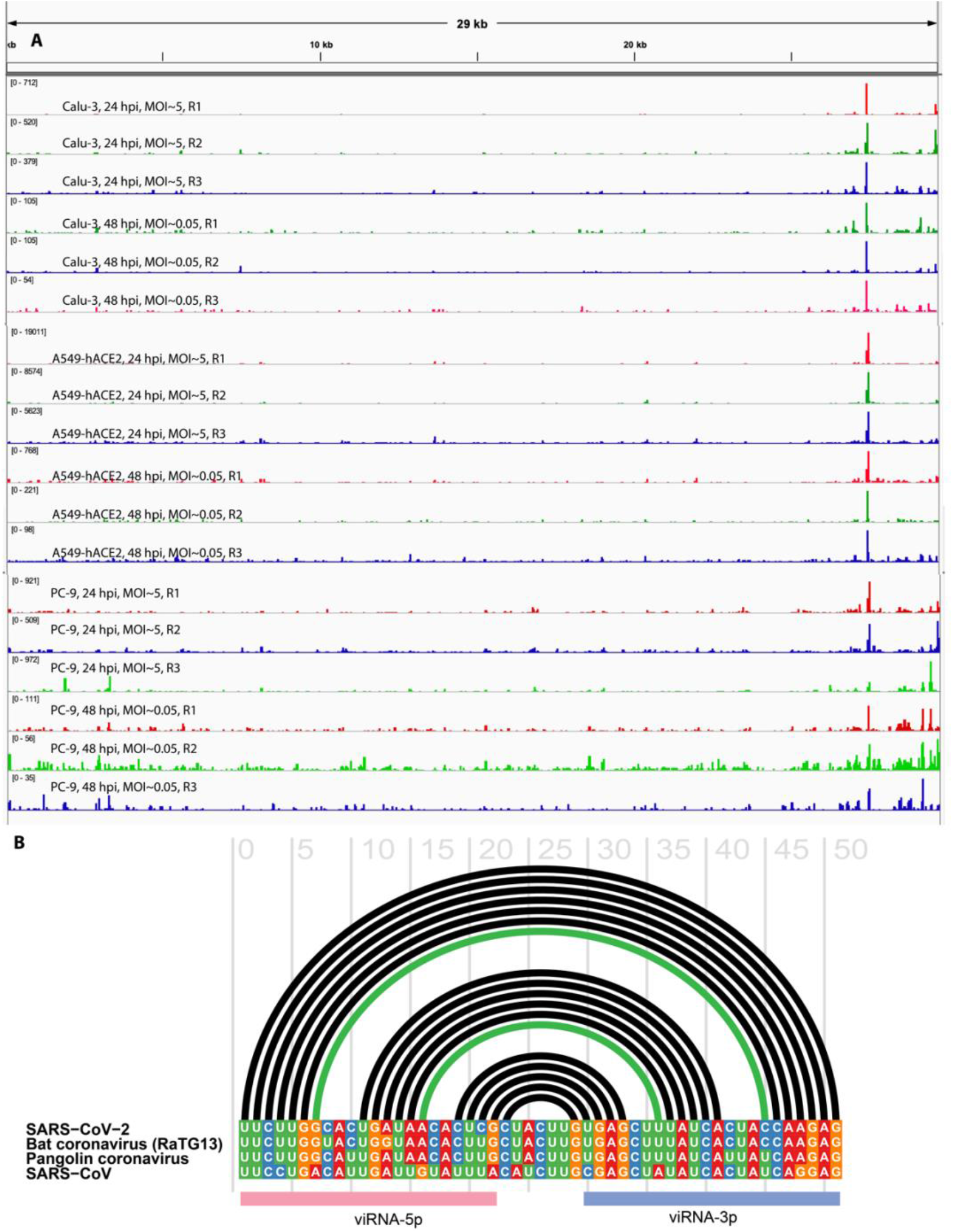
SARS-CoV-2 expresses a small RNA derived from the ORF7a sequence. **A)** Sequencing tracks for all replicates are shown. **B)** Location of vmiR-5p and viRNA-3p annotated on an alignment of coronavirus reference sequences. The hairpin base pairs are indicated by arcs above the sequence alignment with the two previously identified covarying base pairs (21) indicated in green. Alignment and base pairs visualized using the program R-CHI (76).

**Figure S3.**
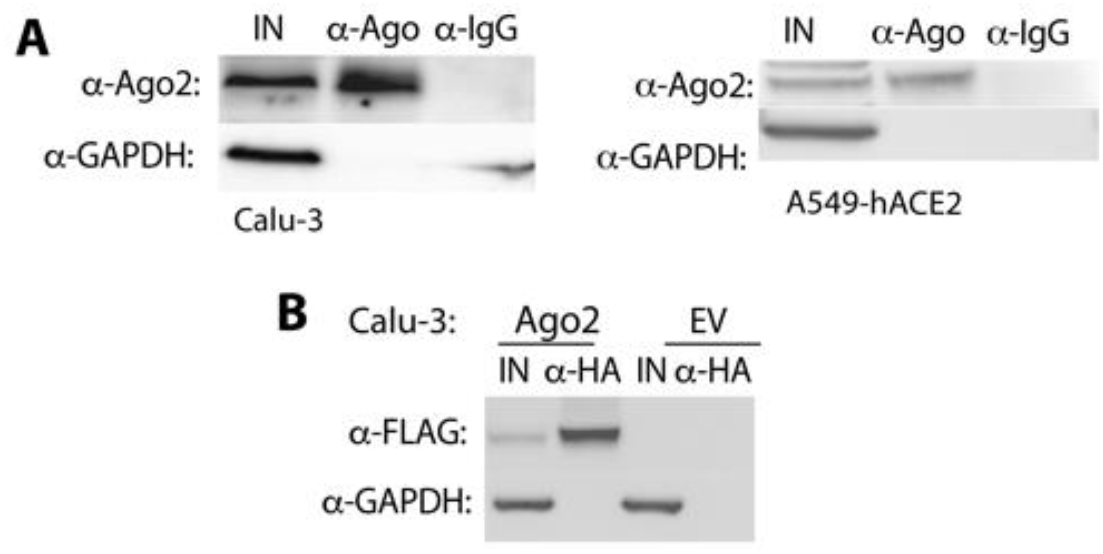
Western blots confirming anti-Argonaute immunoprecipitation. **A)** anti-pan Ago IPs detected with anti-Ago2 antibodies. **B)** anti-HA IP of FLAG-HA-tagged Ago2 detected with anti-FLAG antibodies.

